# Protein Stability Prediction by Fine-tuning a Protein Language Model on a Mega-scale Dataset

**DOI:** 10.1101/2023.11.19.567747

**Authors:** Simon K. S. Chu, Justin B. Siegel

## Abstract

The stability of a protein is crucial to its utility in industrial applications. While engineering campaigns can now be routinely used to enhance protein thermal stability to the level needed in an industrial setting, there is a significant desire to fast-track these efforts through predictive tools allowing one to jump in a minimal number of design iterations to a highly stabilized protein. In this work, we explore utilizing a mega-scale dataset for development of a protein language model tuned for stability. This model is trained on the folding stability of 528k sequences derived from 461 small protein domains and designs, and can accommodate deletions, insertions, and multiple-point mutations. We show that a protein language model can be fine-tuned to predict folding stability. The fine-tuned protein language model, named ESM_therm_, performs reasonably on small protein domains and generalizes to sequences distal from the training set. Lastly, we discuss its limitations when compared to other state-of-the-art methods in generalizing to larger protein scaffolds and highlight the need of large-scale stability measurement on a diverse dataset that represents the distribution of sequence lengths commonly observed in nature.

## Introduction

Protein stability forms one of the foundations of protein engineering and ensuring stability is key in designing resilient proteins. Physics-based methods, including Rosetta [1,2], FoldX [3], and molecular dynamics simulations [4], have found widespread experimental application. Recently the use of machine learning models grounded in biophysical features and evolutionary statistics [5-9], have offered an alternative approach to predicting stability and function without the need for computationally intensive molecular modeling simulations. Fuelled by recent advances in deep learning, convolutional neural networks (CNNs) [10] and graph neural networks (GNNs) [11] are now being adopted to predict mutational impacts on stability, operating directly on the input protein structure [12,13]. For instance, RaSP is a CNN-based model trained on top of Rosetta, a molecular modeling suite [14]. ELASPIC-2, however, operates on both sequence embedding from ESM and structural embedding from ProteinSolver [15-17].

Despite these advancements, the lack of a consistent and universal dataset remains a challenge. While merging smaller datasets into a more comprehensive collection, such as ProtTherm [18], Design to Data [19], and ProtaBank [20], is a feasible approach, the combined dataset are often consisted of closely related but distinct quantities, accompanied by discrepancies in experimental conditions. In parallel, while deep mutagenesis scanning offers profound insights, these studies typically concentrate on a single protein target, limiting the broader applicability of the derived data and models subsequently trained on it. In light of these challenges, Tsuboyama et al. introduced a mega-scale thermostability dataset, encompassing 776k short protein sequences derived from 479 distinct proteins, all consistently evaluated using the same assay [21].

Utilizing this recently developed data set we have fine-tuned a protein language model (pLM), namely ESM_therm_, from ESM-2 [22] to act as an end-to-end stability predictor. We observe that ESM_therm_ performs comparably with state-of-the-art models and can generalize to small protein sequences distal from those in the training set. We also demonstrate that training on an ensemble of protein sequences, instead of mutagenesis studies of a single protein, improves the performance of fine-tuned protein language model for folding stability prediction. Lastly, we discuss the limitations of ESM_therm_ and how it compares with other state-of-the-art methods in generalization to longer protein sequences.

## Results

### Evaluating Model Generalizability on Unobserved Domains

Protein stability prediction can be assessed at different scales of generalizability. Although machine learning algorithms are often trained and tested on different sets of non-overlapping samples, the definition of overlap is more ambiguous for protein sequences. For instance, assigning two point mutants from the same WW domain, one into training set and another one into test set, can assess the model’s generalizability to sequences sharing the same protein domain. However, it does not reflect the model’s generalizability to a completely new domain, such as an SH2 domain. To benchmark our model on both scales, illustrated in Figure 1, our test set is constituted by sequences on protein domains observed in the training set and those only found in the test set.

**Figure 1.**
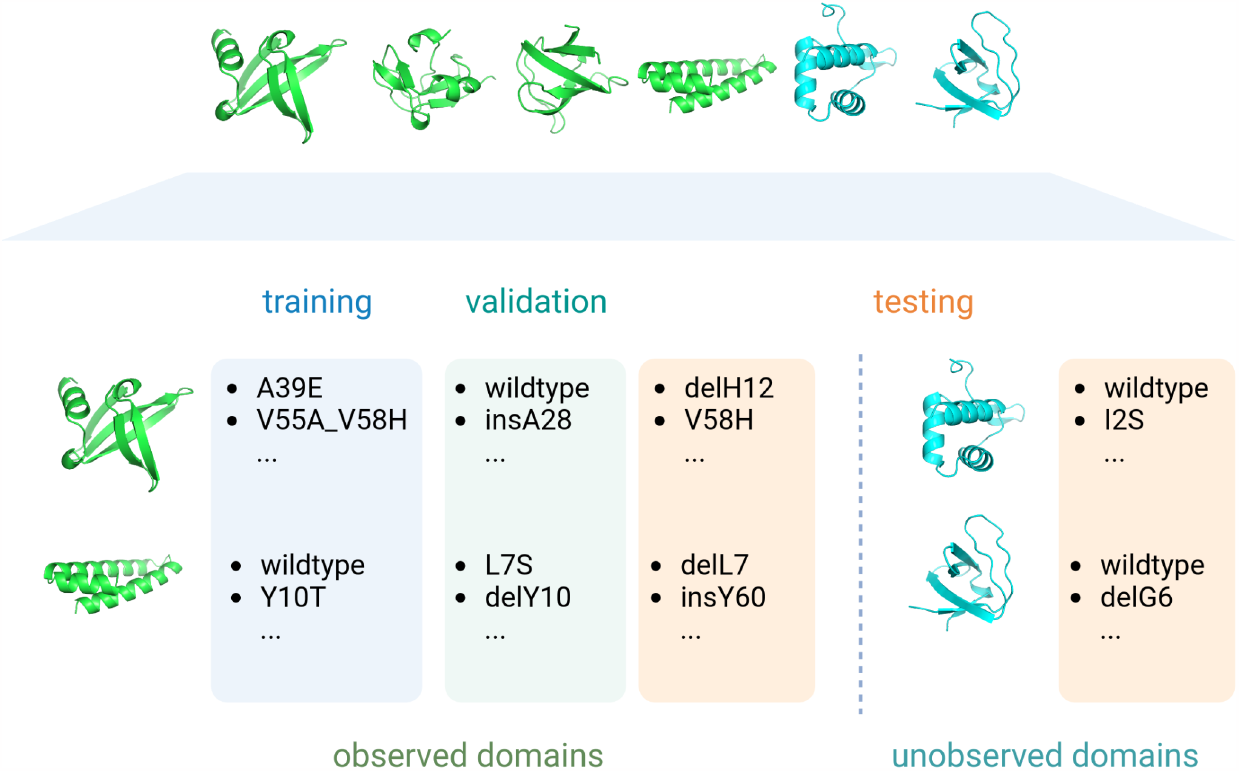
Schematic of dataset preparation. Protein domains are separated into observed domains in green and unobserved domains in cyan. Among each observed domain, mutants are randomly assigned into training, validation, and test sets, whereas for unobserved domains, all mutants are in the test set.

ESM_therm_ generalizes reasonably well to 47 protein domains previously unobserved in the training data, further illustrated in Figure 2. Spearman’s R evaluated by individual domains ranges from 0.2 to 0.9, with the exception of uncharacterized bacterial protein yahO (pdb code: 2MA4) [23]. We characterized the novelty of unobserved domains by the closest percentage identity to any training sequences. Among all test set domains, SH3-subunit of chicken alpha spectrin (pdb code: 6SCW) [24] shares the highest 95.8% identity, and correspondingly a Spearman’s R of 0.88. Sharing 59% identity with its closest training protein domain, *Homo sapien* J-domain protein HSJ1a (pdb code: 2LGW) [25] is relatively novel. Its Spearman’s R is one of the weakest but still retains a value of 0.52.

**Figure 2.**
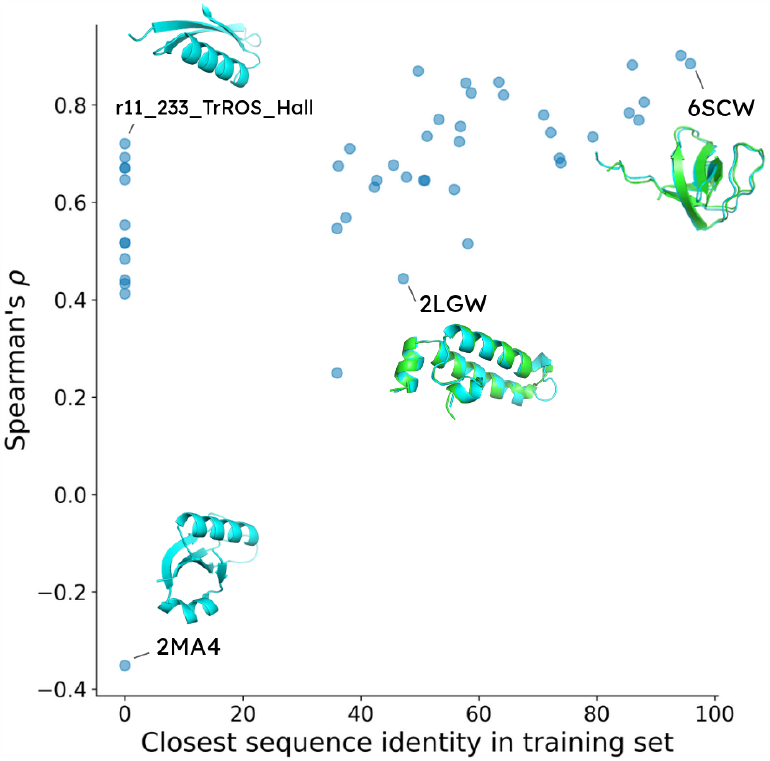
Spearman’s R on unobserved protein domains. x-axis is the closest sequence identity from the evaluated protein domain to those in the training set. y-axis is Spearman’s R evaluated on all sequences from the corresponding domain. The evaluated protein domains from test set are colored in cyan. When homologous sequence is found in the training set, we color the homologous structure in green and align it to the test-set structure.

**Figure 3.**
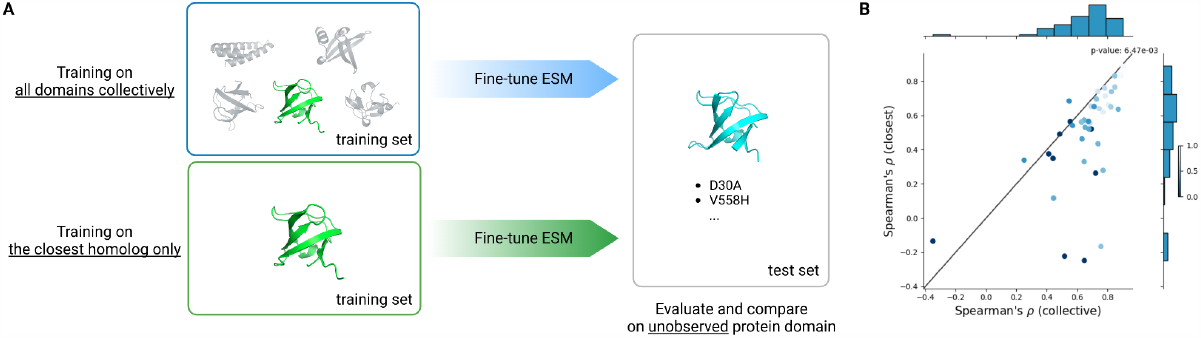
(A) Comparison between transfer learning from homologous protein domain and training on all domains collectively. In the case of transfer learning, we match the test-set protein domain in cyan with the closest sequence homolog found in training set in green by MMseq2 [cite]. (B) Spearman’s R on test-set protein domains absent in the training set. x-axis represents learning from all domains collectively and y-axis is on learning from the closest training-set homolog alone. Domain(s) located under the diagonal line indicates better performance by learning collectively. Homologs were primarily identified by sequence [42], then by structure [30], or discarded when no match was found. Color bar indicates the closest sequence identity to any training-set domains. Statistical significance is performed with Wilcoxon’s rank sum test.

In the 13 cases where no homologous proteins are found in the training set, ESM_therm_ is capable of generalizing to natural and *de novo* designed proteins. For instance, DNA-binding arginine repressor (pdb code: 1AOY) [26] has no homologous training sequences and yet the evaluated Spearman’s R is at 0.69. Similarly, Rosetta-designed αββα domain (HEEH_KT_rd6_0790) and TrRosetta-hallucinated structure (r11_233_TrROS_Hall) [27] have no homologous training sequences. The Spearman’s R on these domains are 0.44 and 0.72 respectively.

### Improving Stability Prediction by Learning All Domains Collectively

Prior to the work by Tsuboyama et al., deep mutagenesis scanning (DMS) is often restricted to a single protein of interest. In the case where the target of interest is not thoroughly mapped, site-saturated mutagenesis studies from a homologous sequence(s) might provide insights into selecting the best mutation(s) for the specific function(s) of interest. However, direct cross-comparison between proteins is often complicated by the difference in measured quantities and experiment conditions between functional assays. This prohibits a systematic assessment on the benefits of learning from multiple target proteins collectively.

Since the folding stability measurements by Tsuboyama et al. were performed in uniform experiment conditions across multiple protein domains, we can now compare between two paradigms, i.e. transfer learning from homologous sequences and learning from all domains collectively, and assess the generalizability of the model fine-tuned on these paradigms on unobserved test-set protein domains.

Extrapolation to unobserved domains benefits clearly from learning all domains collectively. Collective training improves Spearman’s R by on average 0.16 (p-value = 6x10^−3^). CdnL protein (pdb code: 2LQK) [28] shares no sequence identity with any training sequences and was instead matched to its closest structural alignment (pdb code: 2BTT) [29] with Foldseek [30]. Collective training boosted its Spearman’s R from -0.25 to 0.65. Similarly, amino-terminal domain of phase 434 repressor (pdb code: 1R69) [31] was matched by structural alignment and gained 0.74 in Spearman’s R from -0.22. For domains with sequence homologs, WW domain from APBB3 (pdb code: 2YSC) [31] shares 47% identity with its training-set partner (pdb code: 1WR7) [32], and yet still benefits from multi-domain training with an improvement of 0.32. Interestingly, uncharacterized yahO protein remains persistent, where learning on the multiple domains and on the closest homolog scores -0.35 and -0.14 respectively. Overall, this highlights the benefits of a protein stability dataset on a diverse collection of protein domains for the generalization to previously under-studied targets.

Although the uplift brought by collective training highlights the benefits of a consistent large-scale dataset on folding stability, it is still unclear whether the boost originates from the synergy of training collectively or the sheer number of samples. This is especially prominent when the discrepancy in dataset size is significant as an individual domain only constitutes up to 7k sequences, less than 2% of the training set on the collection of protein domains.

### Comparison with Existing Models on Larger Proteins

While natural proteins often span between 200 to 400 residues [33], ESM_therm_ is fine-tuned on sequences no longer than 72 residues in length. To explore its performance under this limitation, we benchmarked our model against six stability-related datasets on larger proteins and compared our results with state-of-the-art covering different methodologies. These include Rosetta Cartesian ddG for molecular modeling, RaSP for structure-based CNN, and ELASPIC-2 which employs a machine learning model based on both structure and sequence embeddings [2,14,15].

We see comparable performance when predicting the thermosability of protein domains absent in the training set. Our pLM achieves a Spearman’s R of 0.65, compared to 0.64 from RaSP and ELASPIC-2 and 0.61 from Rosetta molecular modeling. Drawing an interesting parallel between datasets, Huang et al. reported melting temperatures of beta-glucosidase mutants (pdb code: 2JIE) [19,34] and Romero et al. leveraged a log-enrichment value to gauge the stability for a similar beta-glucosidase (pdb code: 1GNX) [35,36]. Despite their identical alpha-beta barrel scaffold and catalytic mechanics, and a shared 48% sequence identity, the majority of the models achieve a Spearman’s R beyond 0.4 in the Romero et al. dataset, and no method is found to correlate with the Huang et al. dataset. This might be attributed to the difference in sampling methods, in which Huang et al. emphasized mutations around the active site while Romero et al. opted for site-saturation mutagenesis.

Trained specifically on small protein domains, ESM_therm_ does not generalize to other datasets on larger protein sequences. On a collection of direct [37] and indirect [38-40] stability measurements in Table 1, state-of-the-art methods outperform our pLM. Cartesian ddG showcases its generalizability through molecular modeling with a correlation between 0.33 and 0.48. Built on top of Cartesian ddG, RaSP dramatically speeds up the protocol with marginal correlation setbacks. Overall, ELASPIC-2 ranks highest with a Spearman’s R of 0.42-0.58 while our pLM correlates to neither of these datasets.

**Table 1.** Comparison of Spearman’s R across methods on individual DMS datasets. All evaluation is restricted to point mutations, except our pLM on the dataset from Tsuboyama et al. While the mega-scale dataset from Tsuboyama et al. covers multiple protein domains, all other datasets studied on only one target protein.

|  | Rosetta<br>Cartesian<br>ddG | RaSP | ELASPIC-2 | ESM<sub>therm</sub> |
| --- | --- | --- | --- | --- |
| Tsuboyama et al.<br>(test set) | 0.61 | 0.64 | 0.64 | 0.65 |
| Huang et al. | -0.01 | 0.02 | -0.11 | -0.11 |
| Romero et al. | 0.49 | 0.44 | 0.56 | 0.06 |
| Dandage et al. | 0.33 | 0.30 | 0.49 | 0.03 |
| Nutschel et al. | 0.48 | 0.33 | 0.42 | 0.18 |
| Matreyek et al. 2021 | 0.44 | 0.41 | 0.47 | 0.04 |
| Matreyek et al. 2018 | 0.48 | 0.40 | 0.58 | 0.03 |

## Discussion

While our model generalizes reasonably well to new small protein domains in the mega-scale thermostability dataset, it is substantially weaker on larger proteins. Although studies have established a strong correlation between the parallelized assay and direct measurement of thermostability [41,21], we cannot rule out that our language model is biased towards any details specific to this dataset, including experiment conditions and sampling distribution of protein sequences. One hypothesis is that our pLM is biased toward shorter sequences while geometric methods do not suffer from the same pitfall. For instance, the protein domains we trained on are limited to 40 to 72 amino acids in length, a stark contrast to the 177- to 501-residue-long sequences in our additional DMS benchmark. Instead of small protein domains, this might suggest the benefits of a large-scale folding stability dataset on longer sequences for fine-tuned pLM stability predictor.

In parallel, while most methods can rank mutants successfully, predicting the exact value of folding stability is still challenging. Specifically on protein domains absent in the training set, our predictions of absolute folding stability often suffer from an offset and/or scale differently when compared to the experiment. Similarly, other methods might share the same issue. For instance, Rosetta Cartesian ddG follows a different energy unit (Rosetta Energy Unit), and it might not be suitable to be compared directly to kcal mol^-1^. However, the misalignment can be easily resolved by a simple linear regression between model prediction and experiment. Upon recalibration per protein domain, the root mean square error from our model improved from 1.34 to 0.83 and R^2^ from -0.85 to 0.45, averaged across all domains in our test set. For instance, our model scores a negative R^2^ on DNA-binding arginine repressor (pdb code: 1AOY) before recalibration and improves to 0.47 after rescaling, while Spearman’s R remains the same at 0.69 regardless of any rescaling (Figure 4).

**Figure 4.**
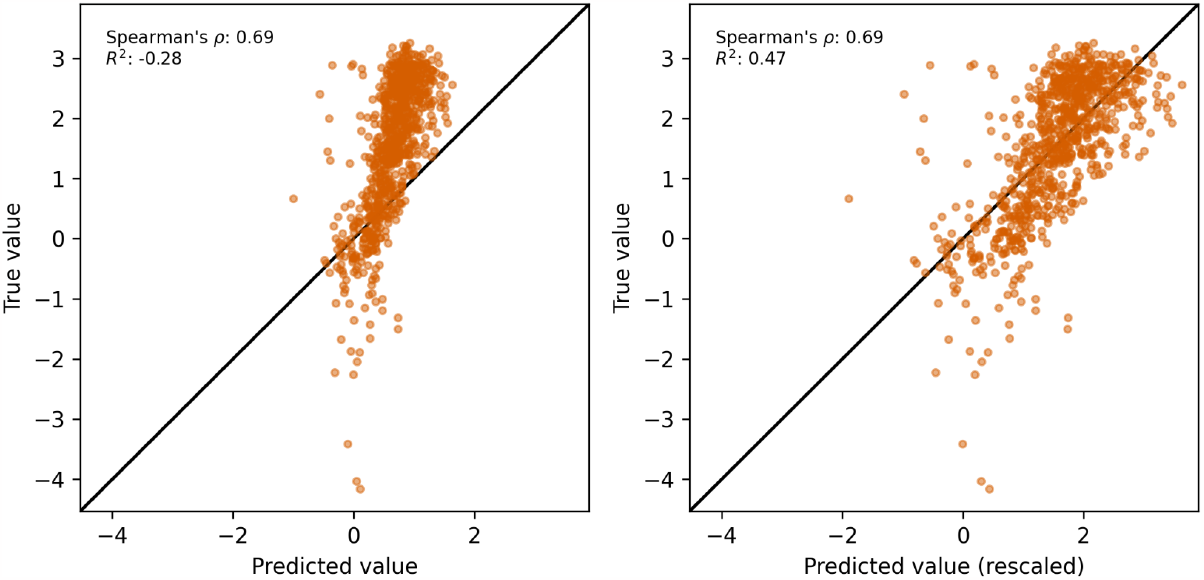
(Left) Miscalibration between prediction and true value on DNA-binding arginine repressor. (Right) Recovered agreement between prediction and true value through linear rescaling on the same set of data.

In this work, we demonstrate that folding stability prediction is possible using a protein language model. Enabled by large-scale protein stability measurements, we fine-tuned ESM-2 on the absolute folding energy of small protein domains. This approach generalizes successfully to protein domains distal from the training set and highlights the benefits of training collectively on all protein sequences instead of mutagenesis study on a single wildtype. While its performance on larger protein scaffolds is lagging behind state-of-the-art, a folding stability dataset beyond small protein domains might be vital to improving the generalizability of a protein language model.

## Methods

### Protein Language Model and Finetuning Protocol

ESM-2 is a transformer pLM pretrained on BERT-task on UniRef50. We finetuned the model on whole-sequence regression task with a classification head on the starting token. We used a local batch size of 128 and a global batch size of 2048. We trained the model on A100 GPU(s) at half precision with patience of 500 steps. We report all test-set evaluations on the checkpoint with the best performance on validation set.

We performed hyperparameter selection on model size (esm2_t6_8M_UR50D, esm2_t12_35M_UR50D, esm2_t30_150M_UR50D, esm2_t33_650M_UR50D) (Table S1), and selected esm2_t12_35M_UR50D to balance prediction performance and compute speed. In addition, we performed an ablation study on pretraining. Model with pretraining has a superior advantage over that with no pretraining (Figure S1).

**Table S1.** Performance evaluation on different model sizes on test set. Metrics are evaluated for each domain/wildtype, and then aggregated into mean and standard deviation over all domains/wildtypes. All models have similar performance metrics with esm2_t12_35M_UR50D except esm2_t6_8M_UR50D on Spearman’s R (p-value < 5x10^−2^).

| Model name | MSE | Spearman's R | R<sup>2</sup> |
| --- | --- | --- | --- |
| esm2_t6_8M_UR50D | 2.48±3.14 | 0.57±0.21 | 0.14±0.23 |
| esm2_t12_35M_UR50D | 2.10±2.52 | 0.65±0.21 | 0.18±0.26 |
| esm2_t30_150M_UR50D | 2.02±2.28 | 0.70±0.15 | 0.15±0.24 |
| esm2_t33_650M_UR50D | 1.91±2.05 | 0.72±0.12 | 0.17±0.26 |

### Dataset Construction

Tsuboyama et al. measured the folding stability of 1.8M measurements derived from 542 protein domains through cDNA display proteolysis. Statistical analysis on distinct DNA but identical protein sequences did not reject the null hypothesis in Kruskal-Wallis test (p-value > 0.99) (Figure S3). We aggregated measurement(s) with the identical protein sequence, regardless of their DNA sequence(s), into a single entry. In cases where the DNA sequence was unique while sharing the same protein sequence, we evaluated the standard deviation of ΔG and log K_50_. We removed measurements when the standard deviation of ΔG is greater than 2 kcal mol^-1^ or that of log K_50_ is greater than 0.5, and kept only domains with at least 100 measurements by protein sequence. This narrowed down the number of entries from 851,552 protein sequences from their original criteria (K50_dG_Dataset1_Dataset2.csv) to 527,785 protein sequences and 416 natural and *de novo* domains.

Under the hierarchical nature of this dataset, by which multiple domains are constituted and each domain holds a collection of multiple mutants, the definition of model generalizability has two layers. The first is the ability of the model to generalize to unobserved mutants on observed domains, and the second is that on unobserved domains. In order to evaluate the model on both observed and unobserved domains, we split our dataset into train, validation, and test sets by domains as illustrated in Figure 1. 10% of all domains are randomly drawn and all their mutants are assigned to test set. Mutants are randomly assigned to train-validation-test sets at an 80-10-10 ratio for the remaining domains.

Instead of training and validating on the same set of protein domains, we attempted to split the protein domains into training, validation, and testing protein domains with no overlap. In other words, the model was finetuned on sequences from training domains, evaluated on unobserved validation domains for early stopping, and tested on unobserved testing domains. We found the model performed worse in this split.

### Comparison between Learning All Domains Collectively and Individually

We finetuned ESM-2 (esm2_t12_35M_UR50D) on each of the 416 protein domains in the training set as our independently learned models. We first matched each test-set domain to its closest partner in the training set by the highest sequence identity by MMseqs2 [42]. In the case where no sequence alignment is identified, we matched test-set domain by the highest structural identity by Foldseek [30]. In the case neither is identified, the test-set domain was not compared. Pairwise comparisons of interpolation and extrapolation are performed in Wilcoxon’s rank sum test.

In addition to extrapolation to unobserved test-set protein domains, a similar comparison on the impact of training on a collection of protein domains was done on interpolation on previously observed protein domains. Overall, interpolation to sequences on observed domains is marginally uplifted by learning from a multi-domain dataset. Illustrated in Figure S2, learning from an ensemble of protein domains weakly outperforms models trained on the same domain by an average of 0.03 (p-value = 2x10^−2^). However, the margin is slim. 72% of the domains have Spearman’s R gain within ±0.1.

### Sequence and Structural Alignment

We implemented sequence clustering and alignment through MMseqs2 [42]. We used MMseqs2 to cluster the domain wildtype sequences hierarchically using a similar strategy in constructing Uniclust database. We dropped prefiltering for all-to-all pairwise alignment by referring to MMseqs2 documentation. For Foldseek [30], we searched the structural identity based on AlphaFold structures from Tsuboyama et al. [21]. Unless otherwise specified, we used the default parameters. The implementation details of clustering and alignment can be found in data/clust/run.sh and src/bertherm/alignment/mmseqs.py under GitHub repository.

### Full-length Protein Benchmark

We identify six stability-related DMS datasets from [43] and another independent mutational (BglB) dataset from Huang et al. [19]. Q59976_STRSQ_Romero_2015 from Romero et al. [36] and BglB datasets share homologous beta-glucosidase sequences but differ in log-enrichment value and melting temperature as indirect and direct thermostability measures. Nutschel et al. reported the thermostability (ΔT_50_) of *Bacillus subtilis* lipase A at different detergent concentrations [37]. Contrary to direct stability measurements, AACC1_PSEAI_Dandage_2018, PTPTEN_HUMAN_Matreyek_2021EN and TPMT_HUMAN_Matreyek_2018 [38-40] correlate with stability through enhancement/depreciation of protein function as an indirect indicator.

### Data and Code Availability

The code and materials are maintained on [GitHub](https://github.com/SimonKitSangChu/EsmTherm). Mutant-level predictions from our model and benchmark evaluation of state-of-the-art are available under Supplementary Materials.

**References**

## Supplementary Materials

**Figure S1.**
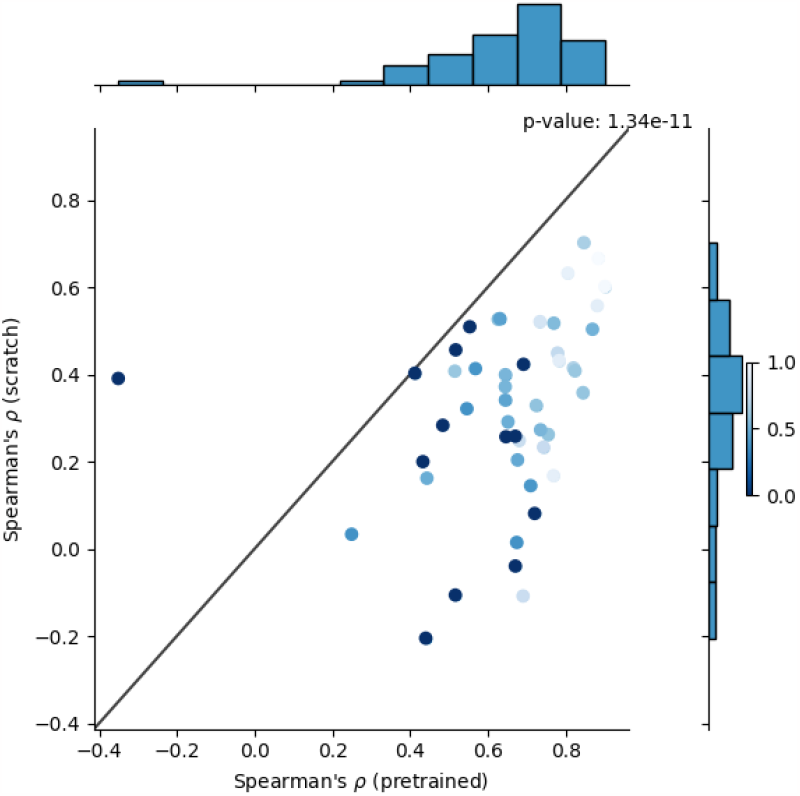
Ablation of pretraining measured in Spearman’s R. Each sample is a collection of mutants of a test-set domain/design. x-axis is Spearman’s R of a domain collection with pretraining. y-axis is that without pretraining. The color bar on the right represents the closest sequence identity in the train and validation set domains. The statistical assessment was performed in Wilcoxon’s rank sum test.

**Figure S2.**
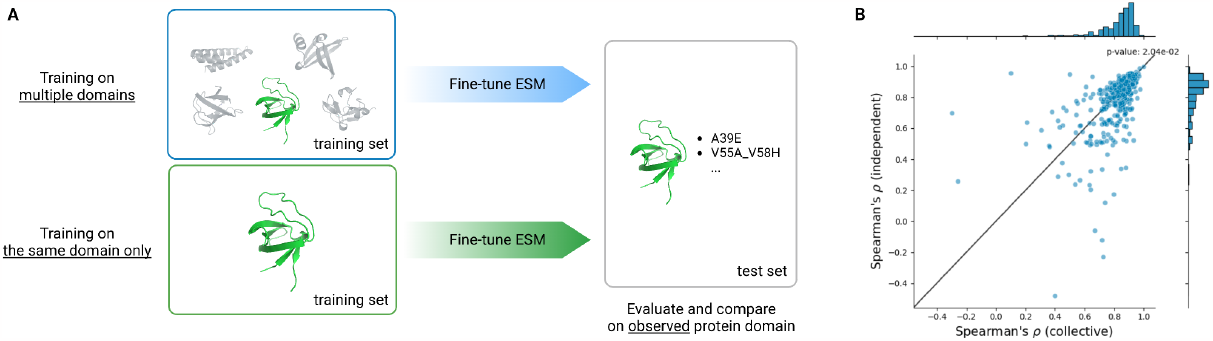
(A) Comparison between transfer learning from the same protein domain only and training on all domains collectively. (B) Spearman’s R on test-set protein domains present in the training set. x-axis represents learning from all domains collectively and y-axis is on learning from the same protein domain alone. Domain(s) located under the diagonal line indicates better performance by learning collectively. Statistical significance is performed with Wilcoxon’s rank sum test.

**Figure S3.**
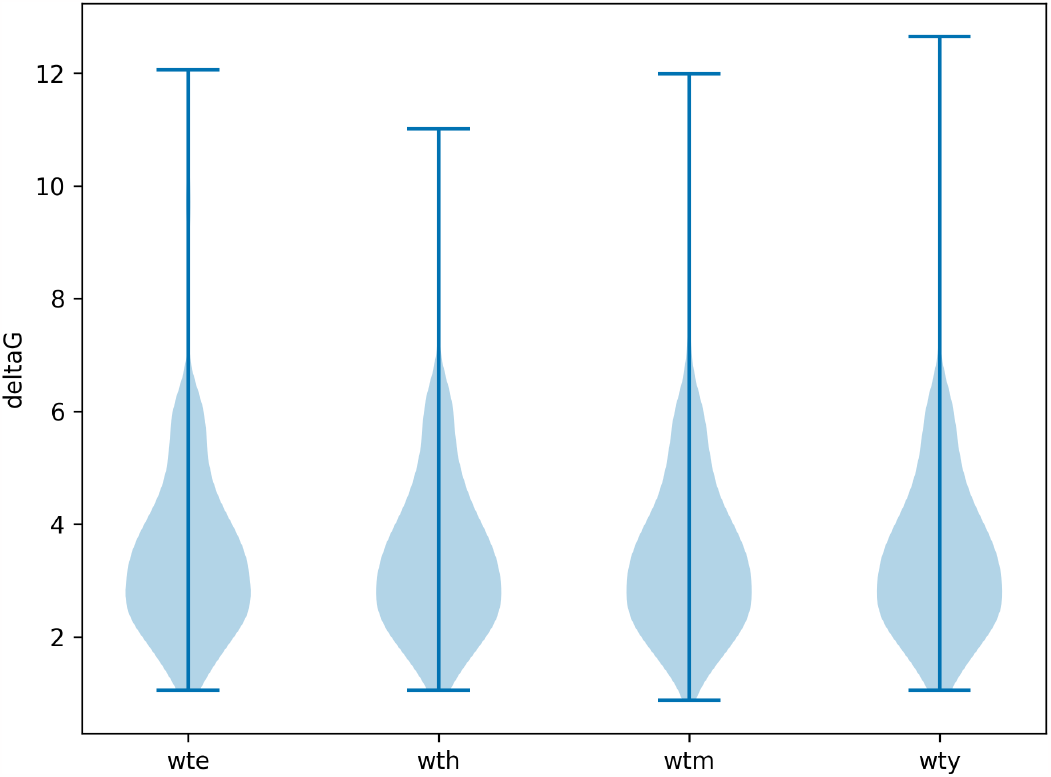
ΔG of homologous sequences (distinct DNA but identical protein sequences) with different wildtype labels. The null hypothesis is not rejected in the KW test (p-value = 1.00).

**Figure S4.**
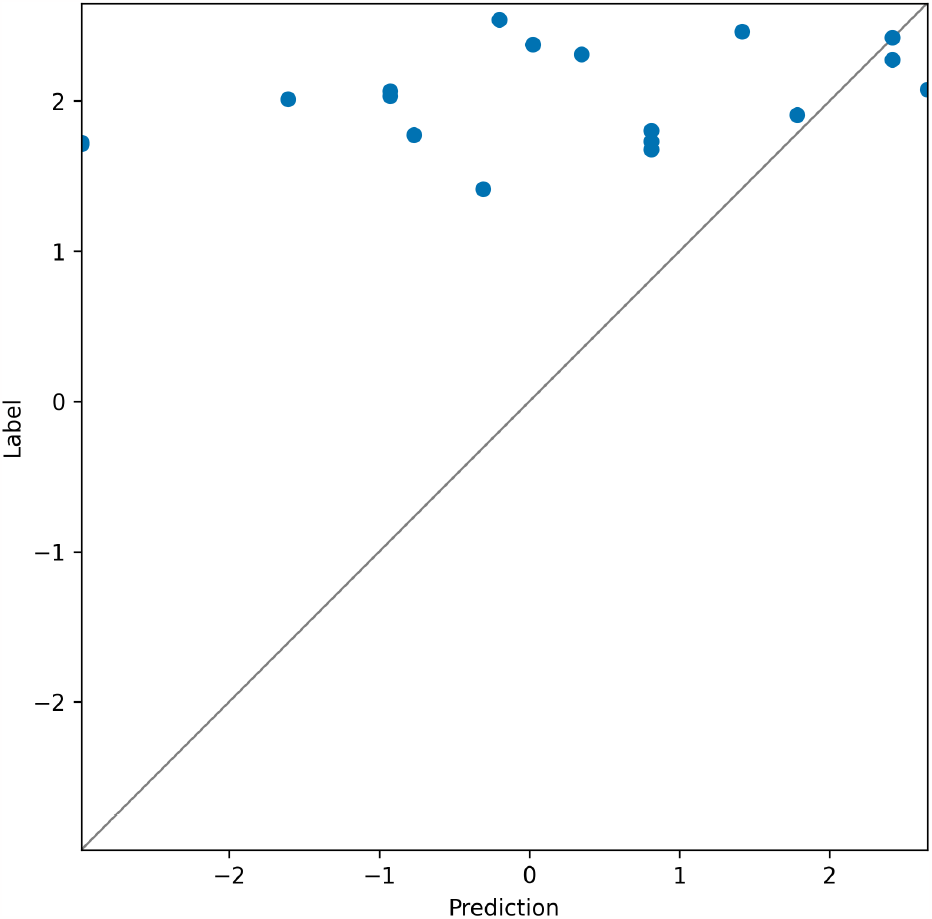
Model performance on predicting ΔG on wildtypes in test set (Spearman’s R = 0.39).

